# 1-methyl-4-phenyl-1,2,3,6-tetrahydropyridine (MPTP)-Treated Adult Zebrafish as a Model for Parkinson’s Disease

**DOI:** 10.1101/2024.06.26.600867

**Authors:** Emmeline Bagwell, Minhyun Shin, Nicole Henkel, Doris Migliaccio, Congyue Peng, Jessica Larsen

## Abstract

Dopamine (DA) is a catecholamine neurotransmitter that works to regulate cognitive functions. Patients affected by Parkinson’s Disease (PD) experience a loss of dopaminergic neurons and downregulated neural DA production. This leads to cognitive and physical decline that is the hallmark of PD for which no cure currently exists.. *Danio rerio*, or zebrafish, have become an increasingly popular disease model used in PD pharmaceutical development. This model still requires extensive development to better characterize which PD features are adequately represented. Furthermore, the great majority of PD zebrafish models have been performed in embryos, which may not be relevant towards age-related human PD. As an improvement, mature *D. rerio* were treated with neurotoxic prodrug 1-methyl-4-phenyl-1,2,3,6-tetrahydropyridine (MPTP) through intraperitoneal injection to induce parkinsonism. Behavioral analysis confirmed disparities in movement between saline-injected control and the MPTP-injected experimental group, with swim distance and speed significantly lowered seven days after MPTP injection. Simultaneously, cognitive decline was apparent in MPTP-injected zebrafish, demonstrated by decreased alternation in a y-maze. RT-qPCR confirmed trends consistent with downregulation in Parkinsonian genetic markers, specifically DA transporter (DAT), MAO-B, PINK1. In summary, mature zebrafish injected with MPTP present with similar movement and cognitive decline as compared to human disease. Despite its benefits, this model does not appear to recapitulate full pathophysiology of the disease with the full profile of expected gene downregulation. Because of this, it is important that researchers looking for pharmacological interventions for PD only use this zebrafish model when targeting the human-relevant PD symptoms and causes that are represented.

## INTRODUCTION

Parkinson’s Disease (PD) is a neurodegenerative condition that affects over 10 million people worldwide, with numbers predicted to increase as scientists continue to investigate^1^. Currently, treatments only improve symptoms, as there is no cure for PD. This is in part due to the difficulty in understanding the etiology of PD, which has been linked to both genetic factors and environmental factors. Several genes are involved in PD, including SNCA, PINK1, PARK2, and LRRK2^2^, with other genes suggested to be implicated still under investigation. Alternatively, pesticides associated with PD include the chemicals glyphosate, paraquat, and rotenone, which can cause oxidative stress which leads to mitochondrial fission and eventual autophagy and apoptosis^3^.

PD is a complex neurodegenerative disorder with several interconnected pathophysiologies, and researchers continue debate which mechanism is responsible for the disease. However, research has identified key factors and mechanisms that contribute PD progression. One hallmark is the progressive loss of dopaminergic neurons in the substantia nigra^4,5^. This loss of neurons leads to a significant reduction in DA levels in the brain, resulting in motor dysfunction and other symptoms. One potential cause of DA neuron loss is the aggregation of protein α-synuclein in the form of Lewy bodies. These protein aggregates are believed to be toxic and disrupt cellular function, leading to neuronal damage and death^6^. Dysfunction in synaptic transmission and plasticity is observed, primarily due to these plaques. The loss of DA neurons disrupts communication within the basal ganglia and between other brain regions, resulting in motor and cognitive impairments^7^. In addition to extracellular protein aggregates, defects in mitochondrial function can result in oxidative stress and energy deficits, both of which contribute to neuronal damage. Typically, mitochondrial fission has been observed in PD patients, leading to mitophagy and eventual apoptosis^8^. The brain is particularly vulnerable to oxidative stress due to its high oxygen consumption. Elevated oxidative stress can damage cellular components, including proteins, lipids, and DNA, and is a contributing factor in PD^9^. Reactive oxygen species (ROS) are also responsible for chronic inflammation within the central nervous system, which is prevalent in PD pathophysiology. Activation of microglia, neural immune cells, can contribute to the release of pro-inflammatory cytokines and further neuronal damage^10^.

All these pathological changes lead to the presentation of PD symptoms, including tremors, loss of motor control, slow muscle twitch, eventual loss of ability to speak, trouble breathing, and freezing episodes of bradykinesia^11^. These symptoms are progressive in nature, though PD has no technical, consistent working timeline, with presentation varying from patient to patient. The field could benefit from the development of improved animal models to assist in both understanding and treating PD. Animal models provide a platform to examine the underlying pathophysiological mechanisms of PD^12^. By introducing specific genetic mutations or exposing animals to toxins or pesticides associated with PD, researchers can investigate how these factors contribute to disease development. However, with the wide range of etiologies, there are many animal models that exist, but none that appear to adequately recapitulate all aspects of human disease^13^. Animals closely related to humans in terms of genetics and neuroanatomy can help researchers better identify common biological principles and mechanisms relevant to PD, which can aid in early diagnosis and monitoring of disease progression^14^.

*Danio rerio*, or zebrafish, have become a promising animal model for PD due to the conserved dopaminergic and catecholaminergic synthesis cascades and neurotransmitter pathways which are implicated in human PD^15,16^, as they have 78% genetic homology to human DNA and age-related changes^17,18^. In addition, high clutch numbers and quick maturity offer large sample sizes, with affordable husbandry as compared to other traditional animal models. By far the most explored zebrafish model for PD is neurotoxin-induced due to its ease of production, hence our focus on 1-methyl-4-phenyl-1,2,3,6-tetrahydropyridine (MPTP) induced PD.

MPTP is a synthetic compound that gained notoriety in the context of PD research due to its ability to induce PD-like symptoms in both humans and animal models^18^. When it is metabolized in the body, it produces a compound called MPP+ (1-methyl-4-phenylpyridinium), which is highly toxic to DA neurons in the substantia nigra, a brain region affected in PD^19^. MPP+ is structurally similar to DA, which allows it to be taken up by DA neurons via DAT^20^. Inside the dopaminergic neurons, MPP+ interferes with the function of mitochondrial Complex I^21^, resulting in an increase in the production of ROS and mimicking PD cascades. Exposure to MPTP or MPP+ can induce motor symptoms and neuropathological changes that mimic PD. However, it’s important to note that MPTP exposure typically leads to rapid and severe Parkinsonian symptoms, which may not fully capture the slower, progressive nature of idiopathic PD in humans^22^. Despite the age-related behavior of PD, a very limited number of papers have focused on using MPTP to create PD in adult zebrafish, likely because embryonic models offer greater ease of handling^23^. Furthermore, adult zebrafish MPTP PD models have not all been created or analyzed for PD-relevant behavior using consistent methods. Therefore, we have developed an MPTP model of PD in adult zebrafish to discover the full capacity of MPTP to model PD in adult zebrafish and to further validate findings of the few relevant papers. In this paper, we developed protocols to create an MPTP PD model in adult zebrafish (sexual maturity ≥ 1 year of age) and explored the ability of this model to recapitulate aspects of human PD.

## MATERIALS AND METHODS

### Zebrafish line

Ubi:Zebrabow was genetically modified by utilizing Cre recombinase with m:Cherry through plasmid transfection^24^. This model was chosen for its applicability in neural tracking. However, in this study, we did not use the full features of the Ubi:Zebrabow model and instead treated these fish as wild type (WT). The use of this model provides promise towards understanding the effects of MPTP on zebrafish neural tissue more deeply in future studies.

### D. rerio husbandry

Fish were reared in a continuous filtration unit using standard light cycles until they reach sexual maturity (6-12 months of age) with 25 fish per tank. Once sexual maturity was reached, fish were kept in tanks of 3-5 fish for MPTP model induction. All fish injected (MPTP (n=20) or saline (n=17)) were isolated from the continuous filtration unit for best practices. They were fed a uniform diet of pellets and flakes to facilitate optical stimulation. Our timeline is presented in Figure 1.

**Figure 1.**
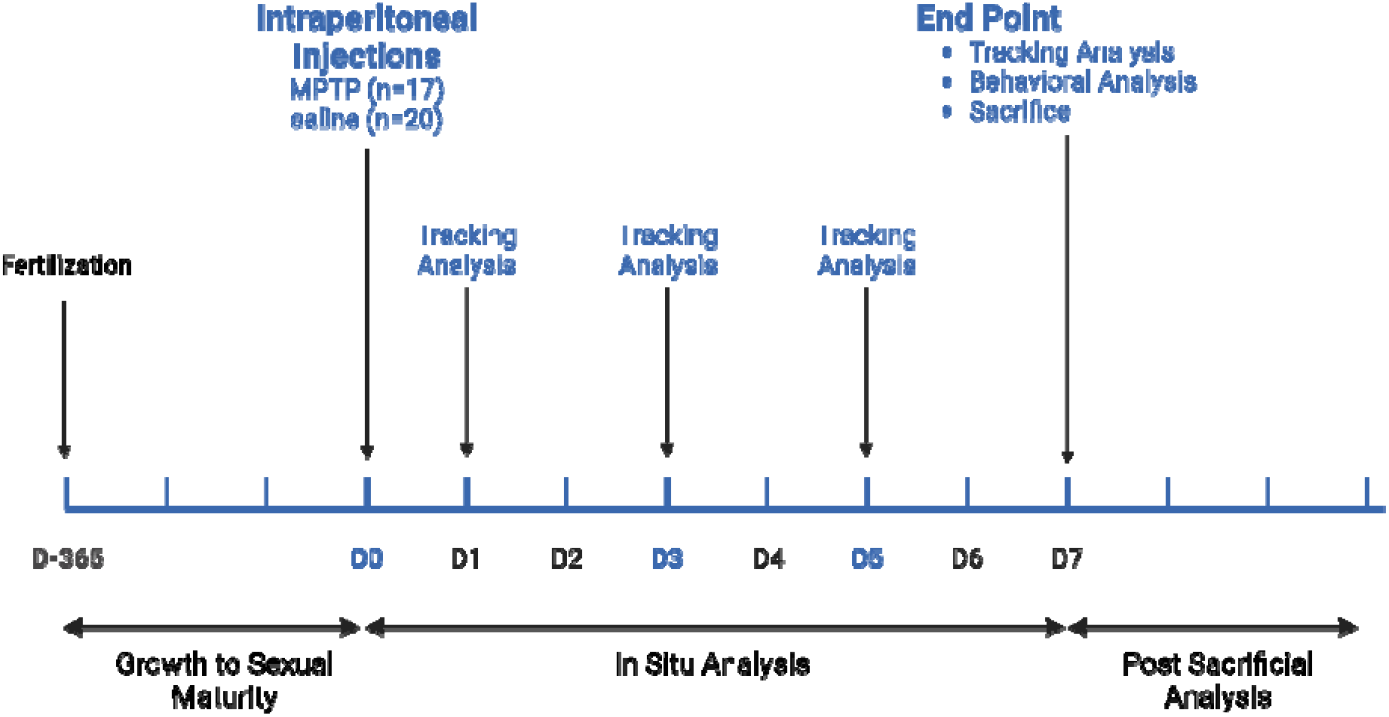
Timeline of fish husbandry and experimental protocols.

### MPTP injections

#### Microneedle pulling

A Flaming/Brown Micropipette puller was used to form microinjection needles (Sutter Instruments P-97). A ramp test was performed to determine the optimal heating temperature to prevent plastic deformation of the needle taper of 492. The pull strength was set to 100, time as 250 milliseconds, with a press at 500 units, and looped 3 times to obtain a tapered needle^25^.

#### MPTP injections

An anesthetizing solution was produced using tricaine mesylate (MS-222) at a concentration of 160 µg/mL utilizing water from the continuous filtration fish rearing tank system (Syndel, Syncaine™). After cutting to hold fish in the dorsoventral position, the sponge was soaked in MS-222 solution and placed in a petri dish for stability. Each fish to be injected was then weighed and an appropriate amount of MPTP injection volume was determined (0.225 g MPTP/1 g body weight). With a working solution concentration of 3.5 µg/mL, the average working injection volume in this experiment was 3.7 µL.

#### Injection Set Up and Intraperitoneal Injections

Injections are performed in the ventral surface, with needle depth extending into the intraperitoneal (IP) cavity. A microinjection system (Microject 1000A BTX Harvard Apparatus) was used to control for uniformity of injection. The injection arm was set to a 45-degree angle from the fish body and the injection time was set to 0.3 seconds with a pressure of 60 kPa (Figure 2). Mature D. rerio were treated either with the neurotoxic prodrug MPTP or volume-matched saline. After injections, fish were placed into tanks with 5 total fish per tank total to recover from injections. Fish were monitored for 30 minutes following injection for potential injury related to anesthesia or injection.

**Figure 2.**
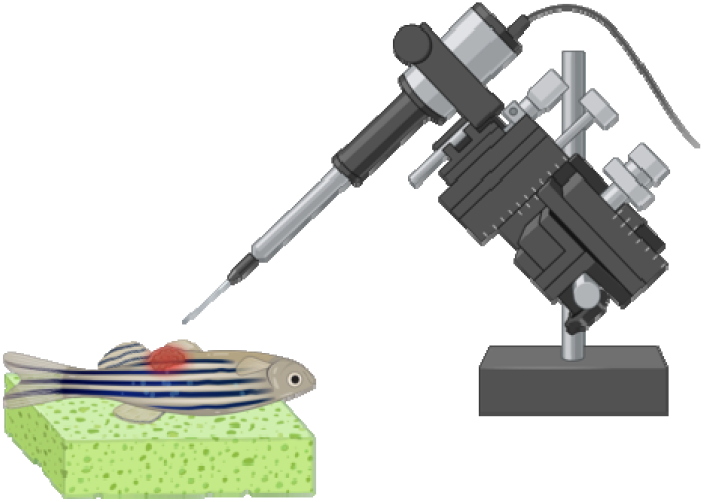
Schematic of the Intraperitoneal Injection Set Up.

**Figure 3.**
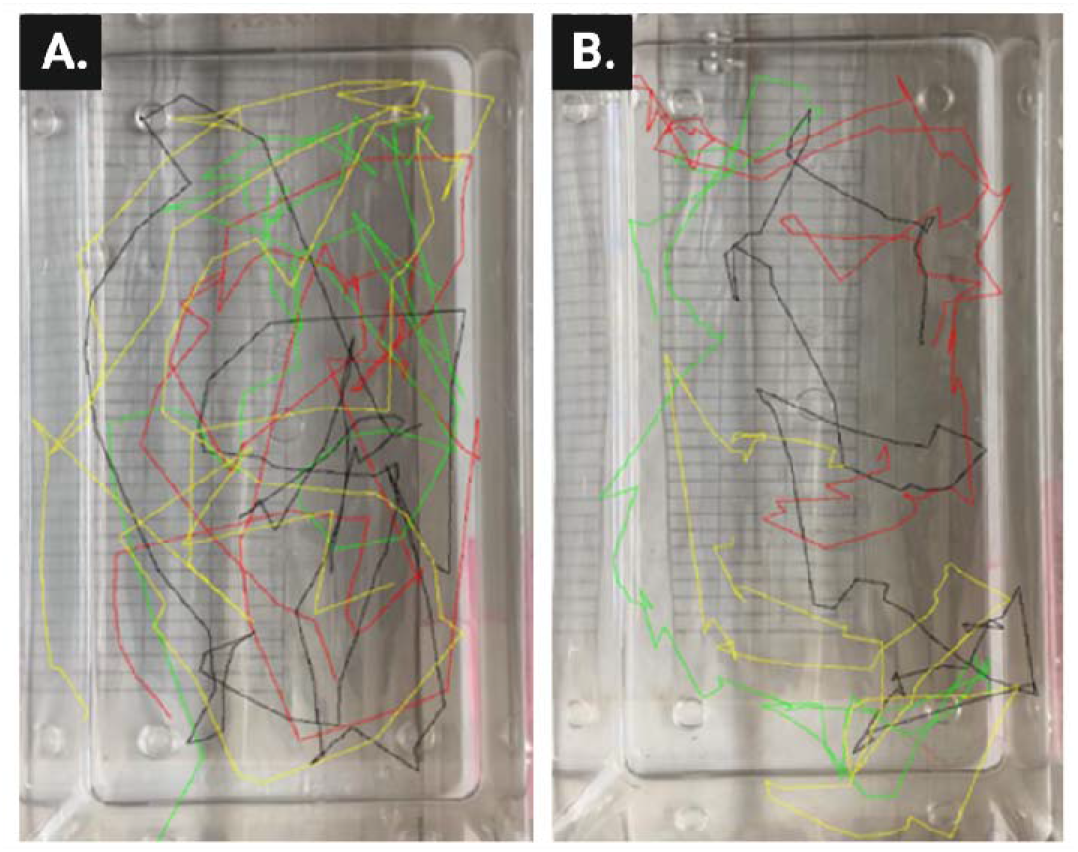
Example motion tracing from Zebtrack on Day 7 post injection. A. Saline injected fish. B. MPTP injected fish. Each tank housed and tracked four fish.

### Motion Tracking

#### Video Set Up

Behavioral analysis through video motion tracking over the course of seven days was performed to confirm the disparity in swim distance between saline-injected control organisms and the MPTP-injected experimental group. An iPhone 13 was placed face-down in a camera stand 8.5 inches above the tank. Fish were allowed to rest for 3 minutes before video was recorded for 1 minute. Video length was modified so that only the middle 30 seconds of video was used in analysis. Each tank was allowed a maximum of 5 fish.

#### Tracking Analysis

To obtain quantitative movement data of zebrafish, an open-source motion tracking program called “ZebTrack” was utilized after optimizations^26,27^. In this program the number of fish in the video, the tracking area, and the dimensions of the fish tank were configured under the “configurations” tab. The processing rate and Threshold were optimized to be set to a value of 5 for each.

### Behavioral Analysis (Y-maze)

A Y-maze was synthesized utilizing a Lulzbot Taz 3D printer and filled with water to serve as a tank for behavioral analysis. Initial studies were done using the control group, followed by an MPTP treated group to avoid absorption of toxic MPTP byproduct water. Fish were placed in the y-maze and allowed 3 minutes to settle prior to a three minute recording. During those three minutes of analysis, entries into arms were noted – with each arm being labeled (Figure 4).

**Figure 4.**
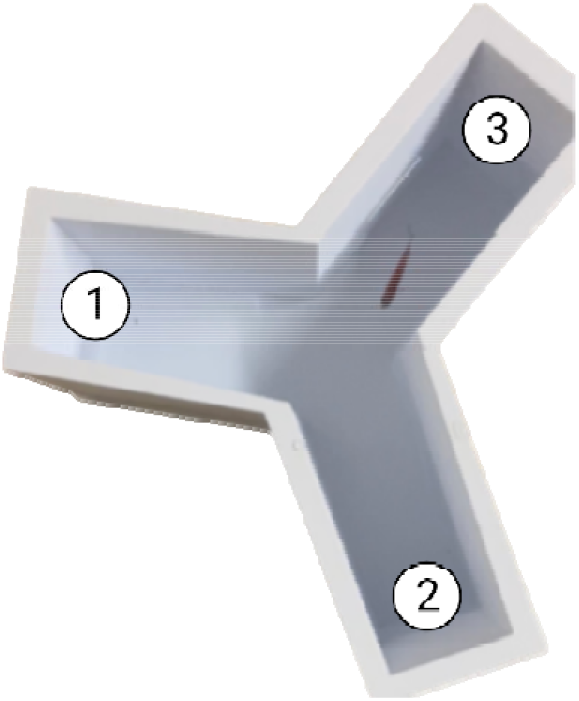
Y-maze used for behavioral analysis of zebrafish. Zebrafish are placed into the y-maze tank. Each arm is labeled with a number (1,2, or 3) and arm entries are recorded for three minutes.

This method takes advantage of a healthy zebrafish’s propensity to explore new areas. If a fish moves into an arm, from any arm, that it wasn’t in previously, that is an alteration and displays good cognitive function. For example, if a fish swims from arm 1 to arm 2 and back to arm 1, this not an alteration or a completion of the maze. However, if a fish swims from arm 3 to arm 1 to arm 2, this represents the completion of a triad (maze completion), an alteration, and good cognitive function. The calculate the alteration percent, we used the equation below ^28^:

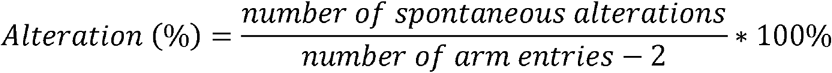

### Postmortem Tissue Preparation

Seven days post injection animals were sacrificed for neural analysis. Tricaine mesylate (MS-222) was prepared in microcentrifuge tubes and stored at -20 °C. Before use, MS-222 was defrosted at room temperature (23 °C) and combined with 300 mL of deionized water to make a euthanizing solution. Fish were placed in the euthanizing solution for 30 minutes. Once euthanized, zebrafish were either stored as heads severed at the operculum or brain tissue for tissue analysis. The head of the fish was removed by cutting sagittally at the operculum. A Leica M890 dissection microscope was used to remove the vitreous matter from the orbital cavity, allowing for visualization of the optic tectum. Using a probe and forceps from a Hamilton Bell Company dissection kit, the skin, foramen of the skull, and pharyngeal jaws were removed to isolate the nervous tissue. Brains were stored in RNA Shield (Zymo Research #R1100-50) at -20 °C.

### RT-qPCR

Nervous tissue stored in RNA Shield was removed from -20 °C and thawed at room temperature. Tissue suspended in RNA Shield prior to being homogenized (Zymo Research #R1100-50). An RNA preparation kit was used to extract purified RNA from the homogenized tissue (Zymo Research #R2052). After using a nanodrop to confirm the concentration of RNA in each sample, a first strand cDNA synthesis kit is utilized to create cDNA from template strand RNA (Invitrogen™ SuperScript™ IV First-Strand Synthesis System, Thermofisher #18091050). The resulting cDNA solution was diluted at a 1:5 ratio for qPCR. A SYBR green probe was selected to add to the qPCR analysis as a fluorescence indicator (iQ SYBR Green Supermix, BioRad #1708880). Primers obtained from Invitrogen (Table 1) are diluted using nuclease-free water to a final concentration of 10µM. Plate preparation includes 5 µL nuclease-free water, 10 µL of iQ SYBR Supermix, 1 µL of forward primer, 1 µL of reverse primer, and 3 µL of cDNA template strand solution. qPCR is run with an initial duration of 10 minutes at 95 °C (once), with the following steps repeating for 40 cycles to determine Ct values of fluorescence in real time: a denaturation setting of 15 seconds at 95 °C, annealing time of 30 seconds at 60 °C, and an extension time of 30 seconds at 72 °C. Due to the low transcript abundance of the genes of interest, we were unable to determine the efficiency of each primer using a serial dilution method. The primer efficiency for each primer set was assumed to be 2. Differential transcript abundance ratios were obtained using a Pfaffl method from the equation below:

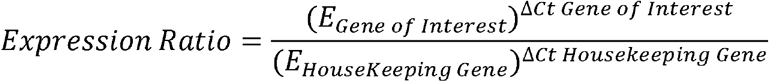

*D. rerio* GAPDH was used as a house keeping gene for result normalization. Each qPCR run was followed by a melting curve scan to monitor the reaction quality.

**Table 1:**
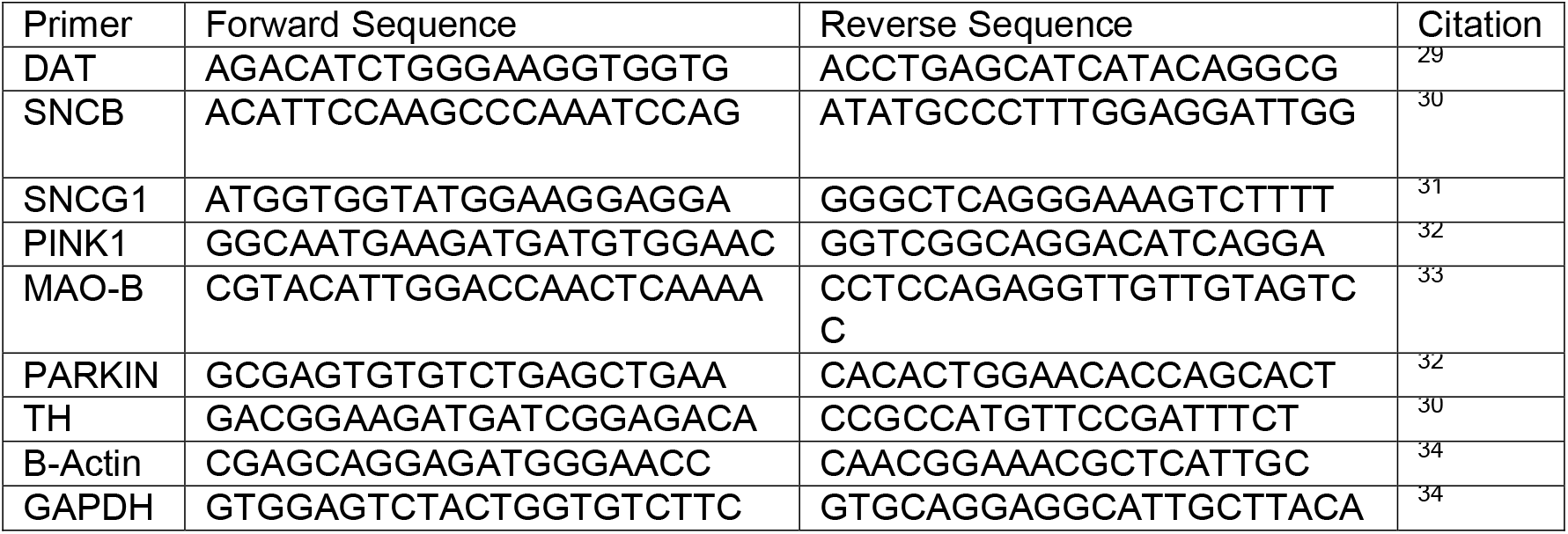
Primers utilized for RT-qPCR.

### Statistical Analysis

Statistically analyses were performed in GraphPad Prism 10.3.1. All data was analyzed confirmed for normality using a Shapiro-Wilk test, due to relatively small sample sizes. Data from motion tracking analysis for distance traveled, speed, freezing time, and alternation were analyzed using an unpaired t-test with Welch’s correction with individualized variance and an α of 0.05, where MPTP-injected fish were compared to PBS-injected fish.

## RESULTS

### Intraperitoneal injections

Both control and MPTP treated groups of zebrafish underwent IP injection to control for any injection-related tissue damage in behavioral and motion analysis. IP injections, in general regardless of treatment group, led to 43.1% mortality in fish, with a mortality rate of 38.5% on Day 1 and 4.6% on Day 2. Of the 65 total fish subject to IP injection, a mortality of 9 control zebrafish and 19 MPTP treated zebrafish occurred because of injection. Fish that survived to Day 3 survived through the end of the study. MPTP, as a neurotoxic chemical, caused a greater mortality rate than saline injections, which was expected.

### Motion Tracking

We observed a significant decrease in average speed and distance traveled of *D. rerio* post IP injection of MPTP in comparison to saline injected controls (equal volume injections) (Figure 5A and B). Specifically, day 1 post injection, speeds of MPTP injected fish declined significantly compared to control from 6.16 ± 0.39 cm/s to 5.33 ± 1.03 respectively. Distances traveled in the analysis window also dropped significantly. Over days 3 and 5 there was no statistical differences observed between treatment groups. However, by day 7 post injection, zebrafish injected with saline traveled an average distance of 78.3 ± 12.7 cm at an average speed of 5.56 ± 0.89 cm/s, which dropped to 43.7 ± 25.1 cm traveled at an average speed of 3.16 ± 1.8 cm/s in MPTP treated fish. Furthermore, on day 7, zebrafish injected with MPTP experienced a significant increase in “freezing time”, indicating that fish were not moving for many seconds during motion tracking analysis (Figure 5C).

**Figure 5.**
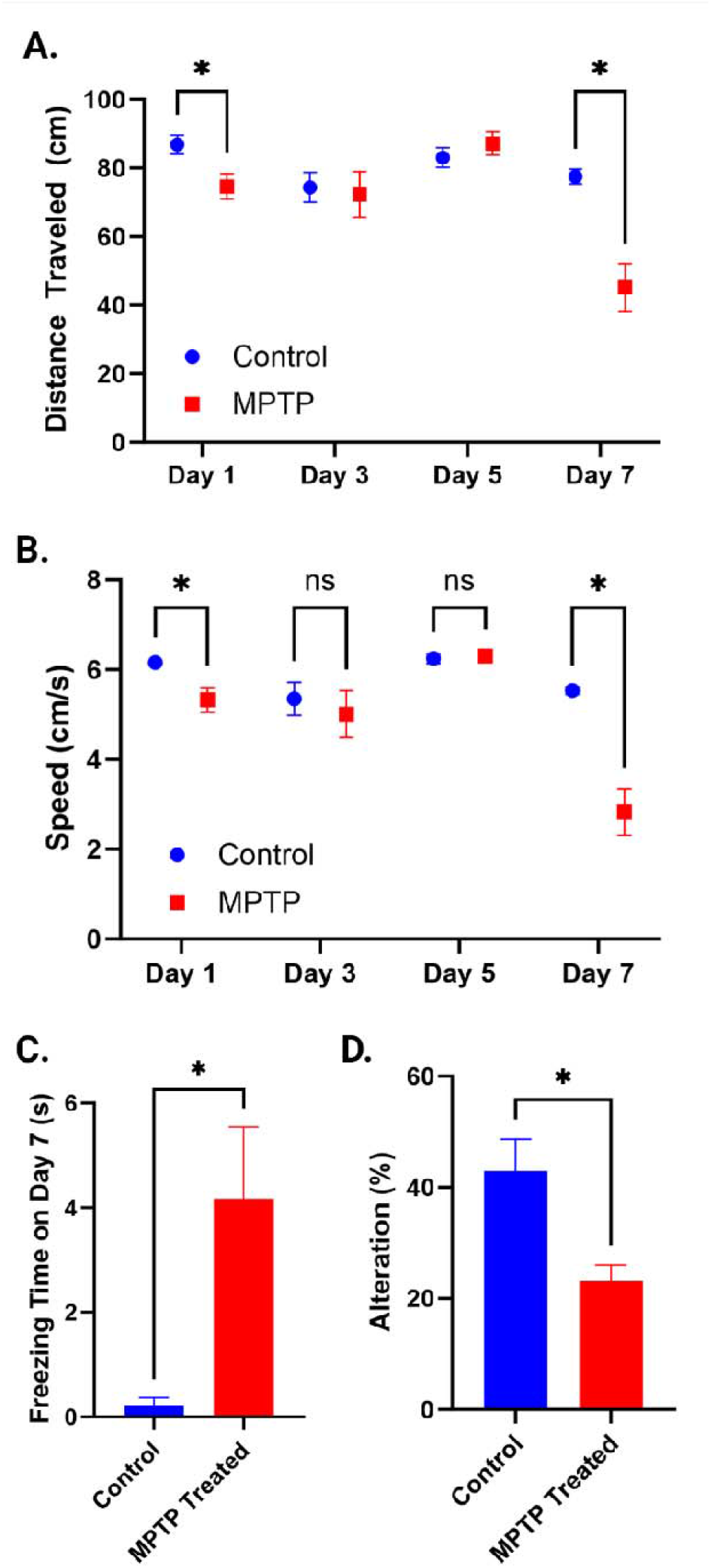
A. Distance Traveled and B. Average Speed as Calculated by Zebtrack in Control (Blue Circle) and MPTP (Red Square) Zebrafish. C. Freezing time for control and MPTP fish at day 7 post injections. D. Alteration of zebrafish injected either with saline (control) or MPTP (MPTP treated) in y-maze analysis. Statistical analysis: *p<0.05

### Behavioral Analysis

MPTP injected zebrafish demonstrated significant levels of cognitive decline compared to saline injected controls. Controls had much higher alteration, at 42.9 ± 20% versus MPTP treated fish with alterations of 23.1 ± 9.3% (Figure 5D).

### RT-qPCR

RT-qPCR assessed any fold changes in gene expression between control and MPTP treated zebrafish (Figure 6). SNCB and SNCG1 were significantly downregulated, with values close to +/-2 being significant. PINK1 is trending towards significant downregulation and TH is trending towards significant upregulation. It appears as if DAT, MAO-B, and PARKIN all remained unaltered with MPTP injections.

**Figure 6.**
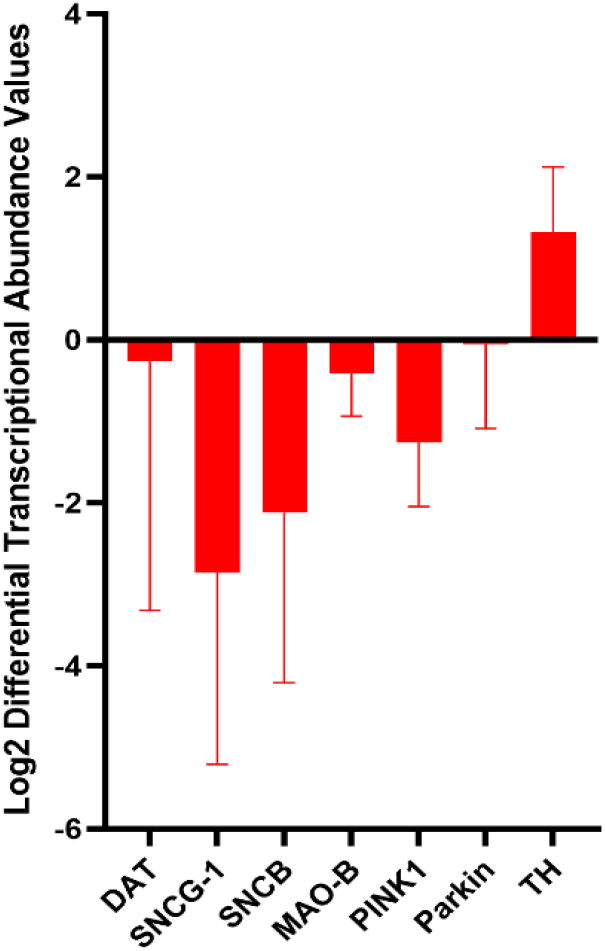
RT-qPCR results of Log2 differential transcript abundance values for PD related genes in MPTP treated fish. Comparisons are drawn to control fish.

## DISCUSSION

With almost identical mortality in MPTP-injected and PBS-injected zebrafish, we believe IP injections led to relatively high mortalities. According to Kinkel, et al., it is possible to perform these IP injections with zero mortality using a 35G beveled needle with a 10 µL NanoFil microsyringe^35^. However, we injected a relatively toxic substance, MPTP, in comparison to their injection of glucose, which is an endogenous sugar. We also have seen that the nature of micropipette pulling can lead to variation in tip sizes and tip roughness, measured via scanning electron microscopy, even with the same overall pulling settings used^36^, which could change the penetrability of the needle into the IP space and change the overall pressure injected despite using the same settings on the microinjection setup. Focus ion beam milling could improve the surface roughness, but micropipette pulling leads to varying diameter needles by nature. It is known that MPTP injections become toxic to adult zebrafish as concentration is increased, with 50% mortality and 100% mortality being observed after cerebroventricular injections of 35- and 100-mM concentrations of MPTP respectively^37^. Because we were injecting at a concentration 13 mM, and based on this literature, we did expect some mortality at less than 50%.

Seven days after one injection of MPTP (0.225 µg/g), we observed statistical decreases in distance traveled and average speed and a statistical increase in freezing time. Distance traveled and average speed dropped by 42 and 49% respectively, while freezing time increased 17.9-fold. This is a similar trend to what was observed in adult zebrafish injected with MPTP intracerebroventricularly, where distance travelled dropped by approximately 47% and freezing activity increased by approximately 51%. However, these fish received four injections before witnessing this behavior^38^. In this regard, IP injected MPTP appears to be a more facile approach to obtaining movement symptoms like those observed in human PD. Another group injected adult wildtype zebrafish with either a single or a double dose of 50 µg of MPTP and found that fish experienced more freezing bouts after only a single injection but did not see a statistical change in either freezing duration or distance traveled compared to control until fish received a double dose^33^. In this case, fish were injected with much less MPTP than in our protocol, which may be why multiple doses were required to obtain PD-relevant movement behavior. Due to the trauma experienced by the fish during and following IP injections, we believe that a single dose is preferred. As a matter of fact, one group performed a single IP injection of MPTP (200 µg per fish) and saw statistical decreases in locomotor activity, similar to what we observed, which provides further validation to this specific approach^39^. The decreased speed (Figure 5B) and freezing behavior observed (Figure 5C) in our fish is very similar to the human PD behavioral symptom of bradykinesia, which presents as a difficulty associated with initiating and maintaining movement and akinesia respectively^40^. It is important to keep in mind that current dopaminergic medications appear to be unhelpful in restoring healthy motor behavior. Therefore, our model may be a useful tool in screening novel pharmacological interventions to assist in restoration of neuromotor function.

Interestingly, we saw some differences in our RT-qPCR results in comparison to the literature. We observed a large amount of heterogeneity among individuals. Like Kalyn and Ekker, we used three brains for RT-qPCR analysis post MPTP injection, although they used a higher concentration of MPTP (25 mM, almost twice our concentration) and cerebroventricular injections for their zebrafish study. They found that whole zebrafish brains had statistically upregulated PARKIN and PINK1 and statistically downregulated DAT, with MAO-B and TH changes being insignificant^38^. Our results were less conclusive with PINK1, PARKIN, DAT, MAO-B, and TH all showing changes that are not statistically significant. These differences observed in gene expression could be due to the decreased bioavailability of MPTP in the brain when injected IP versus cerebroventricular, which is a direct to brain injection. However, human PD patients present with decreased PINK1 protein expression, so the decrease in PINK1 gene expression identified in our study may be more in line with human pathology^41^.

Another study identified that *th1* gene expression dips after IP injection of MPTP in zebrafish, but self-regulates and begins to improve over time, showing that both MPTP concentration and time post injection significantly impact the expression of this gene^42^. We only looked at gene expression on day 7 and therefore the suggested upregulation of TH in our results may be due to the time allowed to pass post injection before analysis. The high individual variation may indicate the post injection *th1* transcription may vary between fish. A future study with more replication will be helpful in analyzing factors that impact high individual variation in post injection *th1* transcription.

Zebrafish do not present with the same synuclein proteins as humans. Instead of producing α-synuclein, zebrafish produce homologues that behave differently, namely β-synuclein (SNCB) and gamma b synuclein (SNCG1). B-synuclein is believed to have a protective role against α-synuclein aggregation in humans ^43^ while gamma b synuclein has been suggested to behave similarly to human α-synuclein^44^. Both related genes were statistically downregulated in our study, which has been linked to decreased motor activity and neural DA levels in other zebrafish studies^45,46^. However, SCNA genes tend to be elevated in human PD^47^, making the value of this model to recapitulate synuclein-related changes unclear.

## CONCLUSION AND FUTURE DIRECTION

In this paper, we demonstrated that an adult zebrafish model using IP injections of MPTP can be useful to assess motion and behavioral deficits associated with PD. However, certain biologically pertinent facets of this model remain to be definitively verified. RT-qPCR results indicate that this disease model may clarify PINK1-related expression changes in human PD. However, there is a lack of clarity in the field demonstrated by papers with conflicted findings on the way MPTP affects these PD related genes. This could be due to differences in MPTP doses and injection routes applied in the adult zebrafish model. It is highly recommended that more in depth studies with much larger zebrafish numbers and varying amounts of MPTP be performed to better understand the genetic and proteomic expression alterations that occur in this model.

While the RT-qPCR results presented here are promising, a western blot analysis would serve to corroborate the downstream effects on translation and the functionality of proteins encoded by specific genes. Nevertheless, conducting western blot experiments in zebrafish presents considerable challenges, primarily due to the limited availability of antibodies tailored for zebrafish and the associated high costs^48^. Furthermore, supplementing the western blot approach with more extensive tissue analysis could clarify the structural changes within cellular and nervous tissues in zebrafish.

Zebrafish have dopaminergic neurons, which makes them useful in modeling neurodegenerative disorders. However, their DA neurons are not organized in the same way as in the human brain. In humans, the loss of dopaminergic neurons in the substantia nigra is a hallmark of PD, leading to motor symptoms^49^. Zebrafish do not have a substantia nigra, so their dopaminergic neurons are distributed differently and often the optic tectum and telencephalon comprise homologous regions^50^. Studies of precisely where dopaminergic neuron loss occurs in this model would strengthen its applicability when studying PD.

Enhancement of this model holds the potential to better mimic the multifaceted mechanisms underlying Parkinson’s disease, although it is important to acknowledge the limitations of this model, which have proven challenging to overcome. The tetraploid nature of zebrafish complicates its direct translation to mammalian organisms (Davis et al., 2014). To validate the findings obtained from zebrafish-based studies, it is strongly advisable to employ a mouse model to corroborate the occurrence of PD in these animal models.

Despite these differences, this model still offers promise for pharmaceutical development and investigation. In addition to the ease of modeling, the results of this study could potentially confirm the use of an adult MPTP model of zebrafish in the development of pharmaceutical products aimed at bradykinesia, confusion, cognitive decline, and other gait-related issues due to its high ability to recapitulate these aspects of cognitive decline. However, in order for this to occur the field will need to garner a deeper understanding of the precise impact of MPTP dosing on zebrafish neurologic changes and develop more uniform procedures for the induction of parkinsonism.

## ACKNOWLEDGEMENTS

The authors would like to extend thanks to Clemson University’s Maker Space for support with material production for this project, specifically the Y-Maze. Additionally, special thanks to the Clemson Aquatic Animal Research Laboratory and John Smink for their support and expertise with zebrafish experiments. All zebrafish experiments were performed under Clemson University IACUC AUP #2021-062. Many thanks to Clemson REDDI lab with resources support from Dr. Delphine Dean, as well as students Mya Beasley and Madison Sexton, for their work and collaboration. Support for this work was partially provided by Clemson University’s Core Incentivized Access R-Initiatives Program.

